# A non-coding indel polymorphism in the *fruitless* gene of *Drosophila melanogaster* exhibits antagonistically pleiotropic fitness effects

**DOI:** 10.1101/2020.12.01.406447

**Authors:** Michael D. Jardine, Filip Ruzicka, Charlotte Diffley, Kevin Fowler, Max Reuter

## Abstract

The amount of genetic variation for fitness within populations tends to exceed that expected under mutation-selection-drift balance. Several mechanisms have been proposed to actively maintain polymorphism and account for this discrepancy, including antagonistic pleiotropy (AP), where allelic variants have opposing effects on different components of fitness. Here we identify a non-coding indel polymorphism in the *fruitless* gene of *Drosophila melanogaster* and measure survival and reproductive components of fitness in males and females of replicate lines carrying one or the other allele. Expressing the fruitless region in a hemizygous state we observe a pattern of AP, with one allele resulting in greater reproductive fitness while the other confers greater survival to adulthood. Different fitness effects were observed in an alternative genetic background, suggesting widespread epistatic effects. Our findings link sequence-level variation at a single locus with complex effects on a range of fitness components, thus helping to explain the maintenance of genetic variation for fitness. Transcription factors, such as *fruitless*, may be prime candidates for targets of balancing selection since they interact with multiple target loci and their associated phenotypic effects.

## Introduction

Genetic variation for fitness provides the raw material for selection and genetic drift to cause genetic evolution of populations [1]. The action of both forces, however, tends to reduce genetic variation. This is particularly relevant in the case of traits that are closely linked to fitness and therefor, by definition, under strong directional selection. The classic explanation for the presence of heritable variation for fitness in populations is mutation-selection-drift balance, where standing variation is maintained at an equilibrium between the generation of new variation by recurrent mutation and its reduction through selection and drift [2,3]. Yet most populations typically harbour considerable amounts of genetic variation for traits and fitness—and more than can be accounted for by mutation-selection-drift balance alone [4]. This discrepancy between theoretical expectation and empirical reality constitutes a central and perennial puzzle in evolutionary biology [4,5].

One possible resolution of this paradox is that fitness variation is actively maintained by balancing selection. Initially popularised by Dobzhansky [6], balancing selection is a force actively maintaining two or more allelic variants at a locus. The active maintenance of polymorphism requires that the selective value of an allele depends on the context in which it finds itself [7,8]. Allelic fitness effects can depend on the genetic context within an individual, as in the case of overdominance [9] or reciprocal sign epistasis [10], or the genetic context in the population, as with negative frequency-dependent selection [11] or variable environmental conditions (fluctuating selection, [12]). In the case of antagonistic selection, polymorphism is maintained because the fitness effect of an allele depend on the sex of the carrier (sexual antagonism, [13,14]), or on an individual’s life history stage (antagonistic pleiotropy, [15]).

Antagonistic pleiotropy (AP) occurs when mutations have a beneficial effect on one fitness component but a deleterious effect on another. Initially conceived in the 1950s [15,16], AP has become a major hypothesis for the evolution of ageing, where mutations that increase fitness early in life are proposed to cause deterioration and increased mortality [15,17]. AP could maintain genetic variation if, for example, one allele confers increased early-life fitness and a shorter lifespan, while the other causes a more even reproductive output over a longer life, with both strategies providing similar long-term fitness pay-offs and greater fitness than an intermediate strategy [18,19]. Despite some empirical evidence of pleiotropic trade-offs [20], modelling has shown that the conditions under which AP generates balancing selection and maintains polymorphism are quite restrictive [18,21–23]. This, combined with relatively few empirical examples of AP in nature, has led to a decline in support for AP as a major contributor to the maintenance of genetic variation for fitness [24].

However, recent theoretical and empirical studies have re-ignited interest in AP as a mechanism generating balancing selection. Models of metapopulation structure in fungi [25] and viability and fertility selection in flowering plants [26] have demonstrated a crucial role of AP in maintaining genetic variation for fitness in wild populations. Similarly, Mérot et al. [24] found that AP in fitness effects and the resulting variation in life-history trade-offs is most likely responsible for the maintenance of an inversion polymorphism in the seaweed fly *Coelopa frigida*. More recent theoretical models have further shown that the conditions required for AP to generate balancing selection are less stringent than initially believed. For example, taking into account sex-specific fitness effects or even small variations in dominance between traits or over time may be enough for AP to generate balancing selection under a wider range of conditions [27]. Furthermore, AP may generate excess fitness variance (relative to unconditionally deleterious mutation-selection balance) by slowing the removal of deleterious variation, rather than maintaining it *per se* [8,27]. Together these developments suggest that the proportion of AP genetic variation (and possibly balanced variation) has been historically under-estimated [4], underscoring the need for further experiments that link sequence-level polymorphism with measurements of different fitness components, ideally in both sexes.

In this study we describe AP fitness effects associated with a polymorphism in a non-coding region of the *fruitless* gene (*fru*) of *D. melanogaster*. The *fru* gene is a key component of the sex-determination cascade and is responsible for sex-specific nervous system development and courtship behaviour [28–30]. In line with its crucial functions, *fru*’s protein coding sequence is conserved across insect taxa [31]. Contrasting with the evolutionary constraint that is evident at the larger phylogenetic level, *fru* also exhibits evidence of positive selection [32]. In line with this evidence for micro-evolutionary dynamics, we identify here a polymorphism within the 5’ non-coding region of the *fru* gene, which consists of a polymorphic indel and associated SNPs that segregate at relatively stable frequencies across worldwide populations of D. melanogaster. Assessing the fitness consequences of the two alleles of this polymorphism, we find that one confers higher reproductive fitness in both sexes, while the other results in greater larval survival and, in some cases, adult longevity. These effects further depend on the genetic background in which the alleles are expressed, which may also contribute to the maintenance of this polymorphism. Our study adds to the growing body of evidence for a reassessment of the role played by antagonistic pleiotropy, and possibly balancing selection, in maintaining individual allele polymorphisms and genetic variation for fitness.

## Methods

### Identification of an indel in a polymorphic region of *fru*

A polymorphic region of *fru* was identified by investigating signatures of balancing selection in population genomic data from two collections of wild flies from Raleigh, US (N=205; [33]) and Zambia (N=197; [34]), based on metrics of genetic diversity (nucleotide diversity, Tajima’s D) and linkage disequilibrium (Kelly’s ZnS) (Supplementary Methods 1). Sanger sequencing of the *fru* region using flies from LHM, a laboratory-adapted North American population of fruit flies [35], revealed that the polymorphic *fru* haplotypes are linked to a 43bp indel (Supplementary Methods 1). As this indel produces a difference in fragment length between PCR products of the two major haplotypes, we designated the two alleles ‘Long’ (L) and ‘Short’ (S).

### Fly culture and husbandry

Unless otherwise stated, flies were maintained on corn-agar-molasses medium with a powdering of live yeast in either vials (8ml of media) or bottles (50ml) in 25°C constant temperature rooms at 50% humidity on a 12:12hr light-dark cycle. When required, flies were collected as virgins, every 0-6 hours post-eclosion until sufficient numbers were obtained. Flies were anaesthetised using a CO_2_ pad for short periods of time and manipulated using a fly aspirator.

### Creation of isogenic lines

We created isogenic lines, which were identical apart from ~1% of the genome including the *fru* locus. Isogenic lines were created through initial identification of LHM individuals carrying the S or L allele, and then backcrossing these into an isogenic Canton-S genetic background (*Df(3R)fru^4-40^)* over seven generations using the pupal phenotype *Tb* as a marker (for full details of crossing scheme, see Supplementary Material 2). We used this approach to generate three independent lines each for the S and L allele.

### Generating focal flies

We performed fitness assays on “focal” flies generated by crossing individuals from the isogenic lines to flies from the *Df(3R)fru^4-40^/TM6B* stock. The resulting individuals carried the *fru* allele (L or S) of a line complemented either by the *Df(3R)fru^4-40^* deficiency (D) or by the TM6B balancer chromosome (B). Since the deleted region of the *Df(3R)fru^4-40^* chromosome extends over the *fru* locus, flies which inherit this chromosome (D) are hemizygous for whichever *fru* allele they inherit. The *fru* polymorphism can therefore be studied in isolation in these D flies. The contrast of allelic fitness effects between flies complemented with the D deficiency or the balancer chromosome allows us to investigate the epistatic effects of the *fru* alleles. The cross to generate focal flies also ensures that line-specific recessive deleterious alleles are partially masked, so as to minimally affect fitness measurements associated with the *fru* alleles.

For each line (S1–3 and L1–3), crosses were performed by setting up replicate vials containing 10 virgin isogenic line females and 10 *Df(3R)fru^4-40^/TM6B* males. These vials were left overnight for the flies to mate. To limit larval densities, we twice transferred flies to fresh vials for 4-hour egg lays (~10am–2pm and ~2–6pm). To establish focal flies carrying the *fru* allele paired with either the D complement (wildtype pupal phenotype) or the B complement (*Tb* pupal phenotype), emerging pupae were sorted into separate vials based on their phenotype. Twelve total line sets were thus established, i.e. lines S1–3 and L1–3 in D or B background, hereafter referred to individually as S1/D, S1/B and so forth, or as S/D, S/B, etc. when referring collectively to all 3 lines carrying a particular allele.

### Fitness assays

#### Female fitness

Focal females were mated to males from their own vial before being placed as triplets at 3 days old into vials containing 1% agar and fed by a capillary tube through the stopper containing a 4:1 yeast to sugar solution (6.5g yeast extract and 1.625g sugar per 100ml) at 25°C and 80% humidity. Triplets were maintained until the focal females were 4–5 days old, with new food supplied daily. Triplets were then transferred to new agar vials (0.8% agar) at ~4pm and allowed to lay eggs for 18 hours. Vials were photographed using webcamSeriesCapture (github.com/groakat/webcamSeriesCapture) software and a Logitech HD Pro webcam C920. We used the machine learning program *QuantiFly* (github.com/dwaithe/quantifly) [36] to count the eggs in each picture. Vials where a female died or where bubbles, debris, or other contaminants caused counting problems were removed from further analysis. Fitness was assayed in 3 experimental blocks.

#### Male fitness

Focal males were reared on standard food in vials of 30 mixed sex flies until 4–5 days old. To assay male fertilisation success, focal males were paired with a competitor male from the *Df(3R)fru^4-40^/TM6B* stock. Pairs of males were held in vials overnight. The next morning a virgin *Df(3R)fru^4-40^/TM6B* female was added to the vial without CO_2_ anaesthesia and the two males competed for mating. The males were allowed to compete for 90mins, thereby maximising the likelihood of a single mating while keeping the rate of double matings negligible. The males were then removed and the female left to lay eggs over a period of several days. Once the larvae pupated, paternity was scored using the pupal phenotype. If all pupae displayed the *Tb* phenotype then paternity was assigned to the competitor (*Df(3R)fru^4-40^/TM6B*) male. If pupae were a mixture of wildtype and *Tb*, paternity was assigned to the focal male. Only vials with >10 pupae were included in further analysis to ensure that wildtype pupae would be observed in cases where the focal male obtained a mating. Male fitness was assayed in 3 experimental blocks.

#### Larval survival, sex ratio and development time

Fifty virgin females from the *fru* isogenic lines and fifty males from the *Df(3R)fru^4-40^/TM6B* line were placed together into egg-laying chambers (~2.5cm diameter, 5cm height) to mate and lay eggs. The floor of these chambers was composed of a grape juice/agar mixture (172ml concentrated grape juice per litre) with a small quantity of yeast as a protein source. After 48 hours, once they had acclimatised to the conditions, the flies were transferred to an identical chamber with the same food source and left for a further 24–30 hours to lay the eggs which would become the “focal” larvae assessed in this assay. Newly hatched, 1^st^ instar larvae were picked and placed in groups of 50 into vials containing standard media and left to develop. Newly formed pupae were removed from the vial and placed into new vials depending on their phenotype (*Tb* or wildtype). For each vial and line, we recorded the number of eclosing flies of each sex, the proportion of surviving larvae, and the sex ratio (once all flies eclosed). Development time were recorded as the number of days from when larvae were placed in the vial until eclosion as an adult.

#### Lifespan

Due to the larger number of flies required for this assay compared to the previous assays, focal flies were generated using a slightly different method. Groups of 100 *fru* allelic line females and 100 *Df(3R)fru^4-40^/TM6B* line males were placed together in an enclosure containing a petri dish filled with corn-agar-molasses medium and left to lay eggs overnight. The next day, small sections of the media, each containing a similar number of eggs, were cut out and placed into individual vials. The eggs were then left to hatch and the larvae to develop. As pupae emerged the flies were separated into vials depending on the pupal phenotype (*Tb* or wildtype). The vials were checked daily until sufficient flies for the experiment eclosed on the same day, which occurred 10 days after eggs were laid. All flies used in the assay were virgins and varied in age by no more than 24 hours. Newly eclosed flies were anaesthetised with CO_2_, separated by sex, and placed in vials in groups of 10. Every other day (Monday, Wednesday, Friday), flies were transferred to a new vial without anaesthesia. The number of dead flies at each transfer was recorded and dead flies removed. If a fly escaped this was recorded and counted in the analysis by censuring. This process was continued until all flies had died.

### Statistical analyses

All statistical analyses were performed in *RStudio* [37]. Mixed effects models were fitted using the package *lme4* [38]. All mixed effects models included the flies’ line ID (S1–3 or L1– 3) as a random variable. If the assay was carried out in multiple blocks, this was also included as a random effect. P-values for each model term were calculated using parametric bootstrapping (package *pbkrtest* [39]) based on 1000 simulations.

Egg count output from the *QuantiFly* program was square root transformed (to achieve better model fitting) and analysed using a linear mixed effects model (LMM) with Guassian error. The model included the *fru* allele (L or S), chromosomal complement (B or D) and their interaction as fixed effect parameters.

Male competitive ability was recorded by scoring paternity (focal vs. competitor male) as a binary response variable. A GLMM (generalised linear mixed effects model) with logit link function and binomial error structure was then fitted for this variable, containing the male’s *fru* allele, its chromosomal complement, and the interaction between the two, as fixed effects. We also included a random block effect in the model; only a limited number of competitive trials could be carried out each day and the assay therefore took place over several days.

Larval survival was measured as the number of adult flies emerging from each vial. An LMM with Gaussian error was applied to the log-transformed number of surviving offspring as a response variable. This produced a better fit according to log-likelihood and AIC than using a GLMM with a Poisson error distribution. The offspring’s *fru* allele and chromosomal complement were included in the model as fixed effects. An additional random variable was added to account for the identity of the vial housing each fly before separation at the pupal stage. Sex ratio was calculated as the number of males divided by the number of females which emerged from each vial and square-root transformed. A Gaussian LMM was applied to these values which included *fru* allele and chromosomal complement as fixed effects and an additional random variable to account for differences between individual vials.

Development time was analysed using a Gaussian LMM including *fru* allele, chromosomal complement, sex and their interactions as fixed effects and larval vial and fly line as random effects. Development time was log-transformed to improve the model fit.

Lifespan data was analysed using Cox proportional hazard models (CPH) from the R package *survival* [40]. A model was constructed including *fru* allele, sex and chromosomal complement as explanatory variables. Significance of model terms was assessed with sequential likelihood ratio tests. Additional models were run with single explanatory variables on either the entire or stratified datasets to estimate hazard ratios for significant model terms. Kaplan-Meier survival curves were fitted using functions from the *survminer* package [41].

## Results

### *fru* polymorphism

Our population genetic analysis revealed that SNP variants in the focal *fru* region investigated here occur at intermediate frequencies in the two distantly related populations studied, Raleigh (US) and Zambia (Supplementary Results 1). Given the perfect linkage between SNP variants and the indel we detect in the LHM population, the two major intermediate-frequency haplotypes can be inferred to include this structural variation worldwide (Supplementary Material 1).

### Reproductive success

863 successful female fecundity trials were performed. There was no effect of the *fru* allele alone on the number of eggs laid (p=0.189; Figure 1). However, there was an effect on fecundity due to the chromosomal complement, with D females laying more eggs than B females (p=0.041; Figure 1). Furthermore, there was a significant allele-by-complement interaction, whereby S/D flies laid more eggs than all other genotypes (p=0.031; Figure 1).

**Figure 1.**
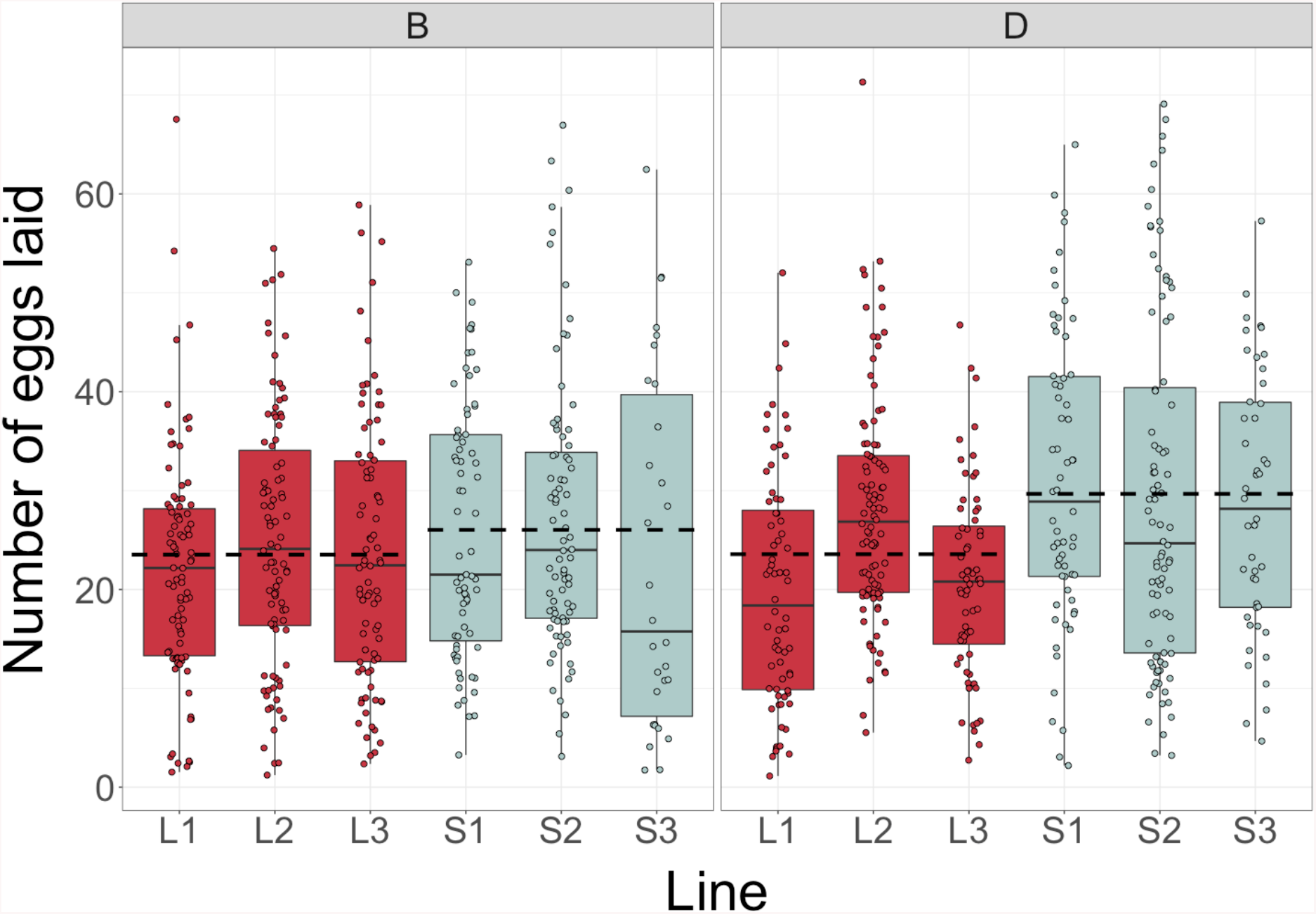
Number of eggs laid by triplets of focal females from each line (L1-3 and S1-3) and chromosomal complement (B and D) over an 18-hour period. Allelic means represented by dashed lines.

We obtained data on mating success for 1149 males. There was no effect of the *fru* allele on male mating success (p=0.562; Figure 2). The success rate of B males was higher than that of D males (p=0.001; Figure 2). S males were particularly good competitors when paired with the D complement, though the allele-by-complement interaction was not statistically significant (p=0.058).

**Figure 2.**
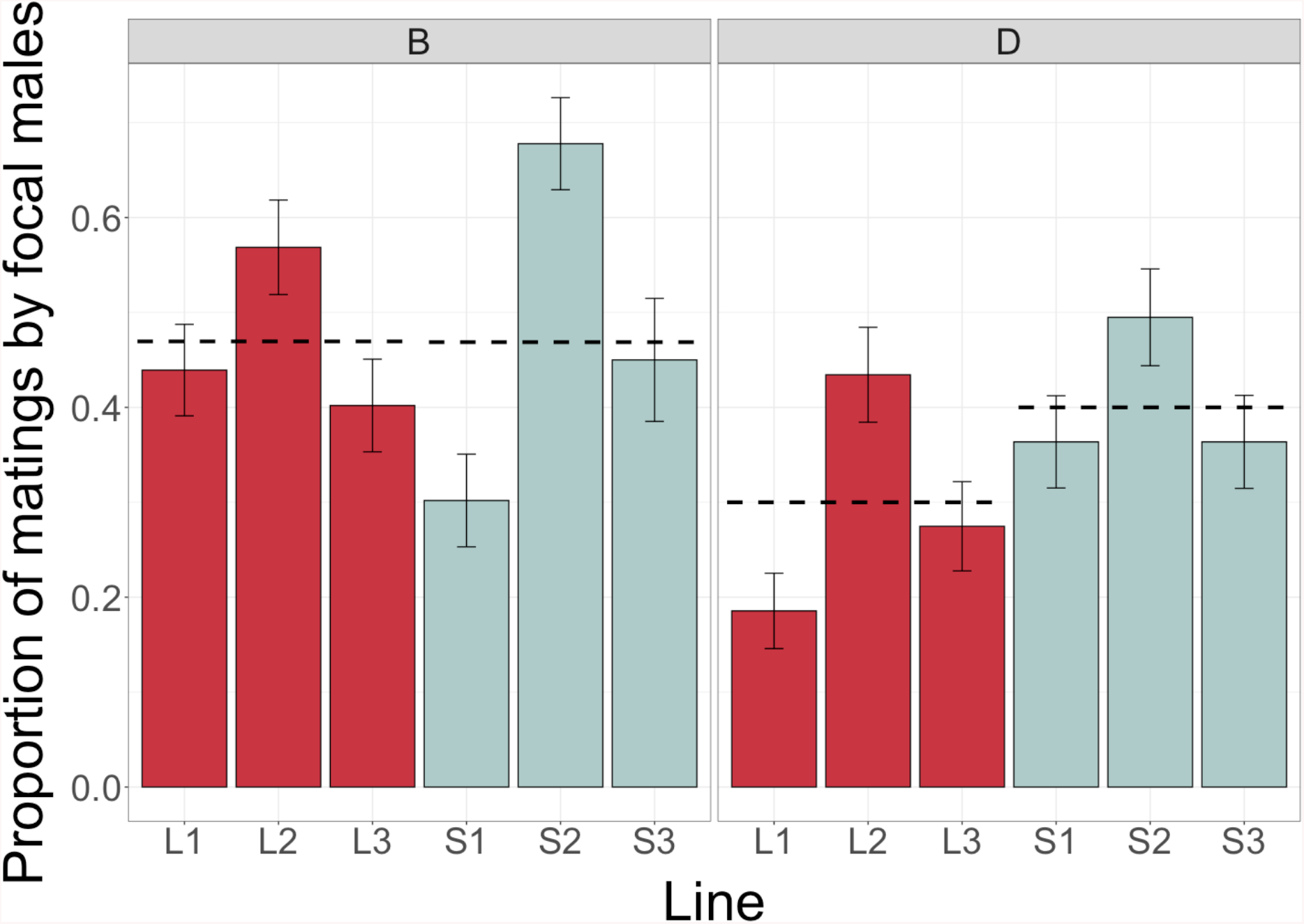
Proportion of matings(±standard error) obtained by focal males for each line (L1-3 and S1-3) and chromosomal complement (B and D). Allelic means represented by dashed lines.

### Larval survival and sex ratio

Data was collected for 2052 flies (1049 females and 1003 males) from 180 vials. A greater number of L allele larvae survived to adulthood compared to S allele larvae (p=0.016; Figure 3) and more larvae that inherited the D chromosome survived to adulthood than those inheriting the balancer chromosome (p<0.001; Figure 3). There was no evidence for an interaction between *fru* allele and chromosomal complement (p=0.275; Figure 3). There were also no significant effects on the sex-ratio of emerging adult flies due to either *fru* allele (p=0.809), chromosomal complement (p=0.158) or their interaction (p=0.097; Supplementary Figure 3).

**Figure 3.**
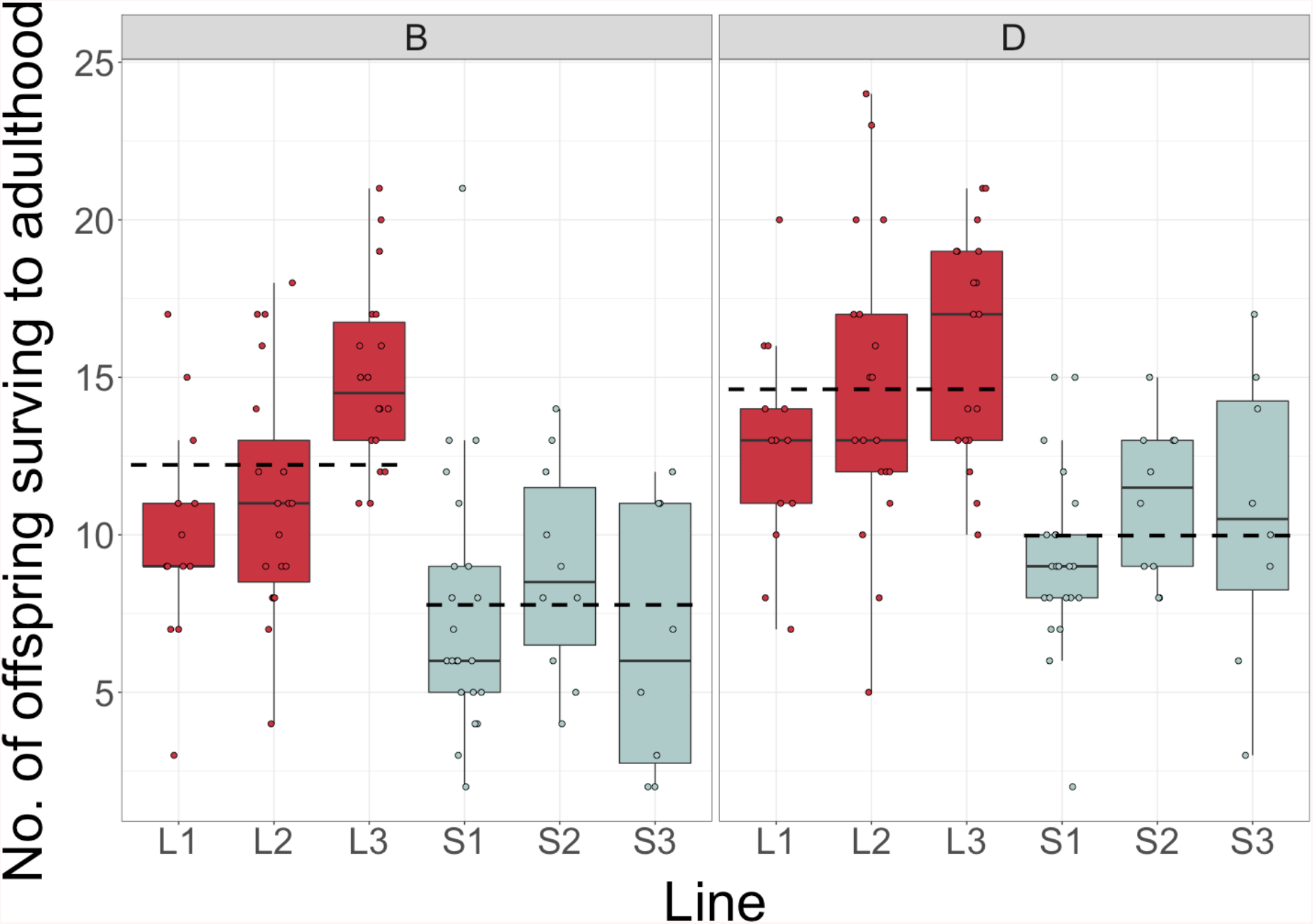
Number of offspring surviving from egg to adulthood for each line (L1-3 and S1-3) and chromosomal complement (B and D). Allelic means represented by dashed lines.

### Development time

Development time data was collected for 2052 flies from 180 vials. Females developed faster than males across all genotypes (p=0.001, Supplementary Figure 4). The *fru* allele had no significant effect on development time (p=0.655). The balancer chromosome lead to faster development than the D chromosome (p=0.003). There was no support for any two-way interactions between these variables (allele-by-sex: p=0.357; allele-by-chromosome: p=0.848; chromosome-by-sex: p=0.106) nor between all three variables (p=0.921) (Supplementary Figure 5).

**Figure 4.**
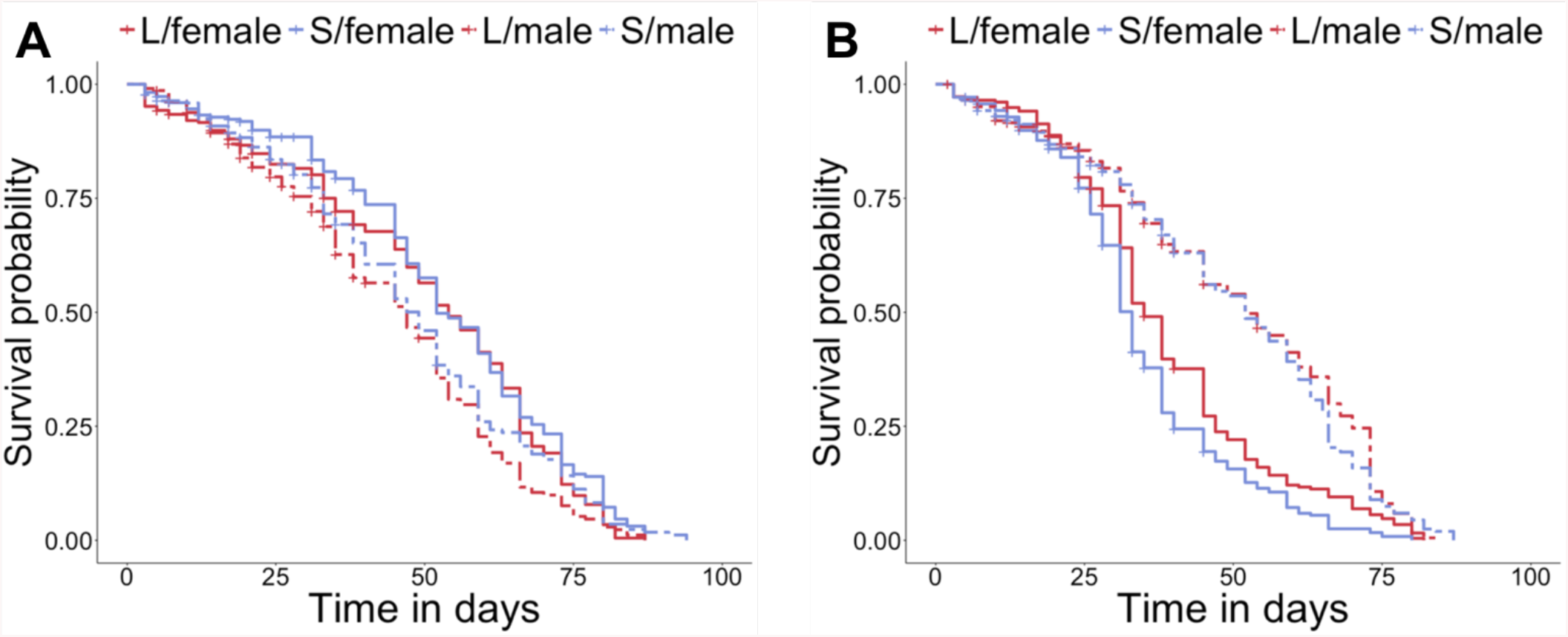
Kaplan-Meier survival curves of flies carrying the D complement **(A)**and B complement **(B)**. Lines represent *fru* allele (L or S, red and blue respectively) and sex cohorts (male or female, dashed or solid lines respectively.

### Lifespan

Complete lifespan data was collected for 1659 flies, with partial data on another 257 flies. A global analyses across the entire dataset did reveal a non-significant effect of allele (p=0.71; Figure 4). We did find, however, a significant effect of complement (p<0.001), with greater lifespan (smaller hazard) in flies with the D than the B complement (HR_D/B_=0.72), and sex (p<0.001), with greater lifespan in males (HR_M/F_=0.82). The latter effect is probably largely driven by a significant complement-by-sex interaction (p<0.001), where the direction of the sex-difference in survival is reversed between the D complement (HR_M/F_=1.27) and the B complement, with a large drop in survival of B females (HR_M/F_=0.50, Figure 4). In addition, we found significant pairwise interactions between allele and complement (p=0.001; D complement: HR_S/L_=0.84; B complement: HR_S/L_=1.14) and between allele and sex (p=0.028; females: HR_S/L_=1.04; males: HR_S/L_=0.93). The three way interaction was not significant (p=0.25).

## Discussion

In this study, we identified an indel polymorphism in the fruitless gene and measured the performance of allelic lines for a number of relevant fitness components. The data provide evidence for complex allelic fitness effects (see Table 1 for a summary), with variation in the impact of the *fru* alleles between fitness components, sexes and chromosomal complements.

**Table 1.** Summary of the effects of *fru* alleles S and L on fitness components, in each sex and for each chromosome complement. The table indicates instances where the S allele or the L allele resulted in significantly greater fitness (S > L and S < L with green and red shading, respectively) and those where no statistically significant difference was observed (S = L, yellow shading). NA denotes cases where a trait could not be measured).

|  | B ♂ | B ♀ | D ♂ | D ♀ |
| --- | --- | --- | --- | --- |
| Female fitness | NA | $S > L$ | NA | $S > L$ |
| Male fitness | $S = L$ | NA | $S > L$ | NA |
| Larval survival | $S < L$ | $S < L$ | $S < L$ | $S < L$ |
| Development time | $S = L$ | $S = L$ | $S = L$ | $S = L$ |
| Lifespan | $S = L$ | $S < L$ | $S = L$ | $S = L$ |

For the cases where the *fru* allele was present in a hemizygous state (paired with the D chromosome) the effects are compatible with AP, in which alleles affect fitness in different and opposing ways (Table 1). Thus, flies inheriting the S allele outperformed L flies in assays of male and female adult reproductive fitness, with S females laying more eggs than L flies and S males tending to have greater competitive fertilisation success. Conversely, flies inheriting the L allele had greater larval survival than those with the S allele in both sexes. These contrasting effects on reproductive fitness and survival suggest that allelic variants at the *fru* locus act antagonistically, contributing to a major life history trade-off.

In addition to AP effects, we also find evidence for epistatic interactions between *fru* alleles and their chromosomal complement. Chromosomes carrying focal *fru* alleles were complemented with one of two paternally inherited third chromosomes, either the wildtype chromosome carrying the deficiency *Df(3R)fru^4-40^* (D) or a balancer chromosome *TM6B* (B). The identity of the complement had additional effects on a number of fitness components, as expected given the large number of sequence differences that will be present between two copies of a major chromosome—in particular deleterious mutations that can accumulate on the non-recombining balancer chromosome. In addition to these additive effects, however, we find that in several cases the fitness outcomes depended on the specific combination of *fru* allele and complement. For example, there was no difference between the effect of the two alleles on adult mortality when paired with the D chromosome, but L flies had lower adult mortality than S flies when paired with the B chromosome. Similarly, S and L males show no difference in competitive ability when paired with the B complement but S outperforms L when paired with D (Table 1). It is important to keep in mind that both complements used in our experiments have genetic particularities (a large deletion in the case of D, a presumably unnaturally high deleterious mutation load in the case of B) that are extreme compared with variation that one would expect to be present in natural populations under to purifying selection. Yet the fact that epistatic allelic differences for particular fitness components arise in the presence of both complements makes it plausible that similar, albeit potentially weaker, effects would occur in interactions of *fru* alleles with naturally occurring polymorphisms elsewhere in the genome.

Life-history traits, such as adult fecundity and survival probability [18,21] that we measured here, are often thought to be associated with genetic trade-offs [19]. In such cases, an increase in performance in one fitness component leads to concurrent decreases in performance in another, for example due to resource allocation. Within this framework, AP is likely to occur when mutations affect the allocation that underlies the trade-off. AP effects have been shown to be able to maintain genetic polymorphism in general models [18,21], models replicating the properties of specific natural systems [25,26] and in empirical observations [24]. Therefore, the antagonistic fitness relationship we have discovered between the two *fru* alleles clearly has the potential to maintain genetic variation at the *fru* locus.

In this context, it is important to note that our findings contradict some of the arguments that had been put forward against a plausible role of AP in maintaining polymorphism through balancing selection [22,23]. Classic theory predicts that in order for AP to maintain polymorphism, fitness effects need to be large and similar across fitness components. This lead to doubts about the ability for AP as a source of balancing selection, based on the assumption that fitness effects are small (≤1%) in most cases [5,22]. Interestingly, however, the fitness differences we observe are considerable. In D flies, where AP is evident, S females lay 25.1% more eggs than L females (29.67 versus 23.57) and S males achieve a third more matings than L males (40% versus 30%), while L flies of both sexes survive to adulthood with a probability that is 46.5% greater than that of S flies (14.62% versus 9.98)%. The efficacy of AP-selection would also be weakened if fitness effects were limited to one sex [22,23]. But this again is not the case here where effects are similar in both sexes for both reproductive fitness and egg-to-adult survival. Reversal of fitness effects between the sexes (sexual antagonism), could have helped maintaining polymorphism in conjunction with AP [27], but does not appear to be present. One property that aids the maintenance of polymorphism via AP and that does not appear in our data is dominance reversal, where the beneficial effect of each allele is dominant (elevated reproductive fitness in both SS and SL individuals, as well as increased egg-to-adult survival in both LL and SL individuals) [23]. If anything, our data suggest that the beneficial effect of the S allele is recessive, and hence only visible when the allele is expressed in the deficiency background. On the other hand, polymorphism at the *fru* polymorphism we study here could be further stabilised by epistasis. Theoretical models don’t often consider epistatic effects in regards to AP but we show through the variable interaction of *fru* with the chromosomal complement that epistatic interactions are present. Epistasis could help maintaining polymorphism if, across a larger set of genetic backgrounds, fitness effects are reversed.

Our results raise the question of how genetic variation at the *fru* locus generates phenotypic effects across the different fitness components we measure. The FRU protein is a BTB-zinc-finger transcription factor and is produced in multiple isoforms, some of which are sex-limited [29,30,42]. The sequence differences between the L and S alleles are upstream of the coding regions, close to the sex-specific promotor P1. Accordingly, the differences observed here between the alleles must arise due to differences in expression levels rather than coding changes, and potentially due to the relative concentrations of different sex-limited and shared isoforms. Both the absolute and relative concentrations of different isoforms could potentially have important consequences on organismal function and phenotypes, given *fru*’s role as a top-level transcription factor. The number of its targets (between 217–291 depending on the particular isoform, [43]) would be expected to generate considerable trickle-down effects through the regulatory cascade. Even slight initial differences in *fru* expression between L and S alleles could potentially result in major, and pleiotropic, effects on a range of phenotypes. For example, mutations in *fru* can result in drastic changes in male mating behaviour and brain development [28,29,44]. The large number of target sites also provides a potential mechanism for the epistatic interactions we observe, depending on the interplay between the abundance of the different FRU isoforms, the specific sites they bind to and the regulation that results from that binding. It is difficult to make inferences about these regulatory effects. But investigation of the sites which interact with fruitless is ongoing [43] and together with a more detailed knowledge of how the target loci are involved in behavioural and morphological traits, this will shed light on the mechanism(s) that link *fru* to downstream traits.

In addition to the effects of allelic variants, complements and their interaction, we observed a significant amount of fitness variation between individual lines carrying a same allele. The method of introgression used to create the allelic lines involved naturally occurring, stochastically placed break points. As a consequence, introgressing a specific allelic variant into the region of interest will also introduce some flanking sequence of unknown size. Variation in the extent of that flanking sequence can generate differences in phenotype between lines carrying an identical allele in the target region. In principle, variation in flanking sequence could also produce systematic differences between S and L lines. In this case, however, the causative variation would require high linkage disequilibrium with the S and L alleles.

Notwithstanding these caveats, our study provides a rare manipulative experimental test of the hypothesis that AP maintains polymorphic variation at an individual candidate gene. Our results provide evidence for allelic variants at the *fru* locus generating a AP relationship between fitness components where one allele (L) enhances survival and the other allele (S) enhances reproduction. Since the *fru* polymorphism influences multiple fitness components, and each allele is beneficial in some instances and deleterious in others, we infer that this polymorphism is maintained through this AP relationship with fitness. Our results complement other recent findings in other systems [24], indicating that AP is a plausible mechanism for maintaining genetic variation for fitness.

## Supporting information

Supplementary methods and results

## Acknowledgements

We are very grateful to Didem Snaith, Harvinder Pawar and Olivia Davidson for their help with pilot experiments, to Florencia Camus for guidance on experimental design and analysis, and to Rebecca Finlay for stock maintenance and media preparation. We further thank members of the MR and A. Pomiankowski research groups for their comments on the results.

