## Supplementary methods and results for "A non-coding indel polymorphism in the *fruitless* gene of *Drosophila melanogaster* exhibits antagonistically pleiotropic fitness effects"

### Supplementary methods 1 – Identification of a polymorphic indel in the *fruitless* gene

Signatures of balancing selection along the *fru* gene

We investigated signatures of balancing selection along the *fru* gene in two wild population samples of *D. melanogaster* flies: a North American population sample of 205 genomes (RAL) and a Zambian population sample of 197 genomes (ZI) [1,2]. Elevated polymorphism and linkage disequilibrium (LD) can both indicate that a given region is under balancing selection [3]. We therefore estimated regional polymorphism (nucleotide diversity, Tajima's D) and regional LD (Kelly's ZnS) over 1000bp windows (500bp step) along the *D. melanogaster* (release 6) genome, in each population, using PopGenome [4].

Sanger sequencing of a candidate *fru* region

A ~1000bp region of the *fru* gene as identified as exhibiting elevated levels of polymorphism and LD in both North American and Zambian population samples (Supplementary Figure 1, Supplementary results 1).To investigate this region in more detail, 96 chromosomes were sampled from LH$_M$, a laboratory-adapted North American population of *D. melanogaster* [5]. Sampling was performed using a 'hemiclonal' approach, in which purpose-built 'clone generator' flies are used to manipulate haploid chromosome sets (X, II, III) [6]. Individual hemiclonal males were crossed with females from a deficiency strain (*Df(3R)BSC509*), which carries a deletion spanning the *fru* gene and a TM6C balancer complement marked with *Stubble* (*Sb*). DNA from the hemiclone/*Df(3R)BSC509* heterozygote offspring of this cross was extracted using standard protocols (see “Phase 1” in Supplementary Figure 2). A ~400bp region of the *fru* gene was then PCR-amplified and Sanger-sequenced using the following primers: 5’-CACCCAACGCCACCTAGTTA-3’ (forward) and 5’-CGCCACTTGATTGCCACATT-3’ (reverse).

**Supplementary Results 1 - Identification of a polymorphic indel in the *fruitless* gene**

We found that a 1000bp-window of the *fru* gene exhibited unusually high levels of polymorphism and local LD relative to the genome-wide average (red dashed line in Supplementary Figure 1). This was true both in the RAL population (upper 2$^{nd}$ percentile of nucleotide diversity; upper 12$^{th}$ percentile of Tajima’s D; upper 5$^{th}$ percentile of Kelly’s ZnS), and in the ZI population (upper 5$^{th}$ percentile of nucleotide diversity; upper 11$^{th}$ percentile of Tajima’s D; upper 9$^{th}$ percentile of Kelly’s ZnS). Sanger sequencing further revealed that this polymorphic region of *fru* segregates for a 43bp indel, producing fragment length differences between the PCR products of the two alternative haplotypes in this region. We designated these haplotypes ‘Long’ (L) and ‘Short’ (S), respectively.

To infer the frequency of the *fru* indel polymorphism in the RAL and ZI populations in the absence of direct indel polymorphism data, we examined the frequency of SNPs located in very close proximity to (<80bp), and in tight LD with, the indel (Supplementary Figure 1). A haplotype network constructed from these SNPs showed that they are found at intermediate frequencies and cluster into allelic classes rather than by population (Supplementary Figure 1). Given the large evolutionary distances between the RAL and ZI populations used in the construction of the haplotype network, this is suggestive evidence that the *fru* indel (and/or alleles linked to it) are under some form of antagonistic and/or balancing selection. We therefore performed further experiments to test this hypothesis.

**Supplementary methods 2 – creation of isogenic lines**

To assess the sex-specific fitness effects of the L and S alleles, we created fly lines homozygous for each allele but otherwise isogenic for a Canton-S background across the rest of their genome ('isogenic allelic lines'; see Supplementary Figure 2 for the full crossing scheme).

First, we randomly selected three lines carrying the S allele and three lines carrying the L allele among the 96 sequenced hemiclonal lines (see Supplementary Methods 1, "Sanger sequencing of a candidate *fru* region") and introgressed these alleles into an isogenic background, as described below. Introgression of the *fru* allele was performed with the help of a *Df(3R)fru*$^{4\text{-}40}$/TM6B deficiency stock, carrying a deletion spanning the *fru* locus (see Supplementary Figure 1) in a Canton-S background, complemented with the third-chromosome balancer TM6B marked with the dominant mutation *Tubby* (*Tb*). Introgression of the *fru* allele onto the deficiency chromosome and into the Canton-S background was achieved by repeatedly backcrossing: (i) females heterozygous for a third chromosome carrying a focal *fru* allele (*fru*$^{S/L}$) and the *Df(3R)fru*$^{4\text{-}40}$ deficiency (themselves obtained by mating the hemiclonal line and females from the *Df(3R)fru*$^{4\text{-}40}$ deficiency stock), with (ii) males from *Df(3R)fru*$^{4\text{-}40}$ deficiency stock (see Supplementary Figure 2). Since balancer and deficiency chromosomes are lethal in homozygous state and balancers carry the dominant *Tb* marker, the wild-type offspring of a hemiclone/*Df(3R)fru*$^{4\text{-}40}$ x *Df(3R)fru*$^{4\text{-}40}$/TM6B cross are always identifiable as *fru*$^{S/L}$/*Df(3R)fru*$^{4\text{-}40}$ heterozygotes. By repeatedly backcrossing *fru*$^{S/L}$/*Df(3R)fru*$^{4\text{-}40}$ heterozygote females to *Df(3R)fru*$^{4\text{-}40}$/TM6B males, the original hemiclonal genome carrying the focal *fru* allele is gradually eroded through recombination in females and replaced with the isogenic Canton-S background of the *Df(3R)fru*$^{4\text{-}40}$ deficiency line. After 7 generations of backcrossing, the allelic lines should carry on average less than 1% of the original hemiclonal haplotype (i.e. 1% of the original X-II-III complement).

Having introgressed the *fru* allele into the Canton-S background of *Df(3R)fru*$^{4\text{-}40}$, we created lines homozygous for the *fru* allele (as opposed to *fru*$^{S/L}$/*Df(3R)fru*$^{4\text{-}40}$ heterozygotes). Because *fru*$^{S/L}$/*Df(3R)fru*$^{4\text{-}40}$ heterozygotes and *fru*$^{S/L}$/ *fru*$^{S/L}$ homozygotes are phenotypically indistinguishable, this was achieved through a two-step crossing procedure. An initial cross served to identify pairs of parents in which both individuals carried a focal *fru* allele. Virgin *Tb*-carrying offspring of a *fru*$^{S/L}$/*Df(3R)fru*$^{4\text{-}40}$ x *Df(3R)fru*$^{4\text{-}40}$/TM6B cross (either *fru*$^{S/L}$/TM6B or *Df(3R)fru*$^{4\text{-}40}$/TM6B) were set up in pairs (dyads A, B, C, see "Phase 3" in Supplementary

Figure 2). Depending on the genotypes of the F1 pair, this cross can either produce: (i) 100% *Tb* F2s, if both F1 parents were *Df(3R)fru*$^{4\text{-}40}$/TM6B—these were discarded, or (ii) some fraction of non-*Tb* F2s, if the F1 pair were *fru*$^{S/L}$/TM6B+*Df(3R)fru*$^{4\text{-}40}$/TM6B or *fru*$^{S/L}$/TM6B+*fru*$^{S/L}$/TM6B. To distinguish the two latter cases and identify pairs of *fru*$^{S/L}$/TM6B individuals that are capable of producing the *fru*$^{S/L}$/ *fru*$^{S/L}$ individuals we required, an additional 'test cross' was performed where F2s were backcrossed to *Df(3R)fru*$^{4\text{-}40}$/TM6 males. Based on the F3 phenotype, the genotype of the F2 could be inferred, as *fru*$^{S/L}$/ *fru*$^{SL/}$ F2s produce a 1:1 ratio of wild-type to *Tb* F3s, whereas *fru*$^{S/L}$/*Df(3R)fru*$^{4\text{-}40}$ heterozygotes produce 1:2 ratio of wild-type to *Tb* F3s. F2s producing a ratio of wild-type to *Tb* F3s that was significantly less than 1:2 (as assessed from a $\chi^2$ test; Supplementary Table. 2) were used to establish isogenic allelic lines.

**Supplementary Figure 1.** Population genetic signatures of elevated polymorphism in the *fru* gene. **A.** Map of the *fru* gene, including breakpoints of chromosome bands, gene model, approximate span of the *Df(3R)fru*$^{4\text{-}40}$ deletion, nucleotide diversity (in RAL) in 1000bp windows (grey horizontal lines = median genome-wide nucleotide diversity; dark grey horizontal lines = 95% quantile of genome-wide nucleotide diversity) and position of the *fru* indel (vertical red dashed line). Alignments of a subset of the ~400bp region spanning the *fru* indel (brackets) obtained through Sanger sequencing of $LH_M$-derived chromosomes are also shown, with closely linked SNPs (used to construct the haplotype network shown in C.) shown as red arrows. **B.** Histograms of nucleotide diversity, Tajima's D and Kelly's ZnS for all 1000bp windows across the genome in RAL and ZI populations, with the vertical red dashed line representing the 1000bp window encompassing the *fru* indel. **C.** Haplotype network constructed from SNPs closely linked to the *fru* indel (red arrows in A.) in RAL and ZI populations.

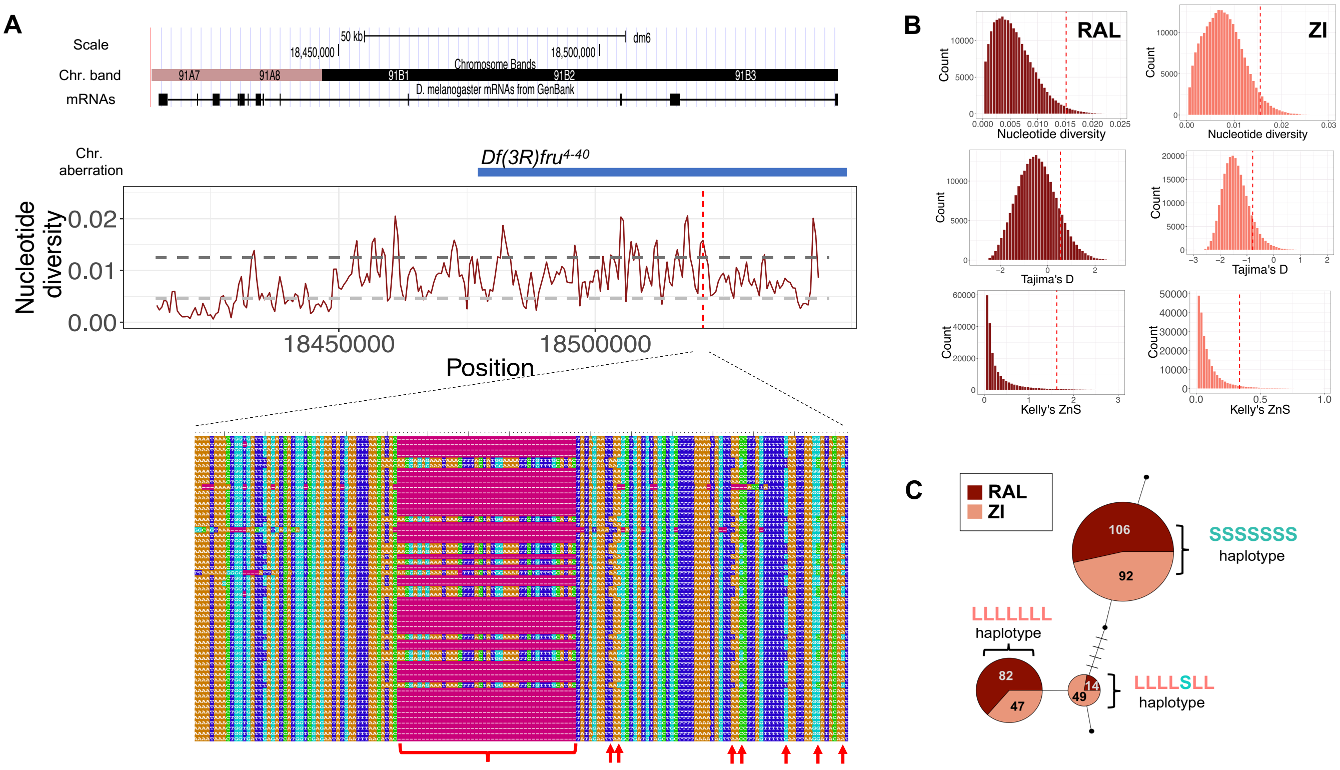

**Supplementary Figure 2.** Crossing scheme used to create isogenic lines.

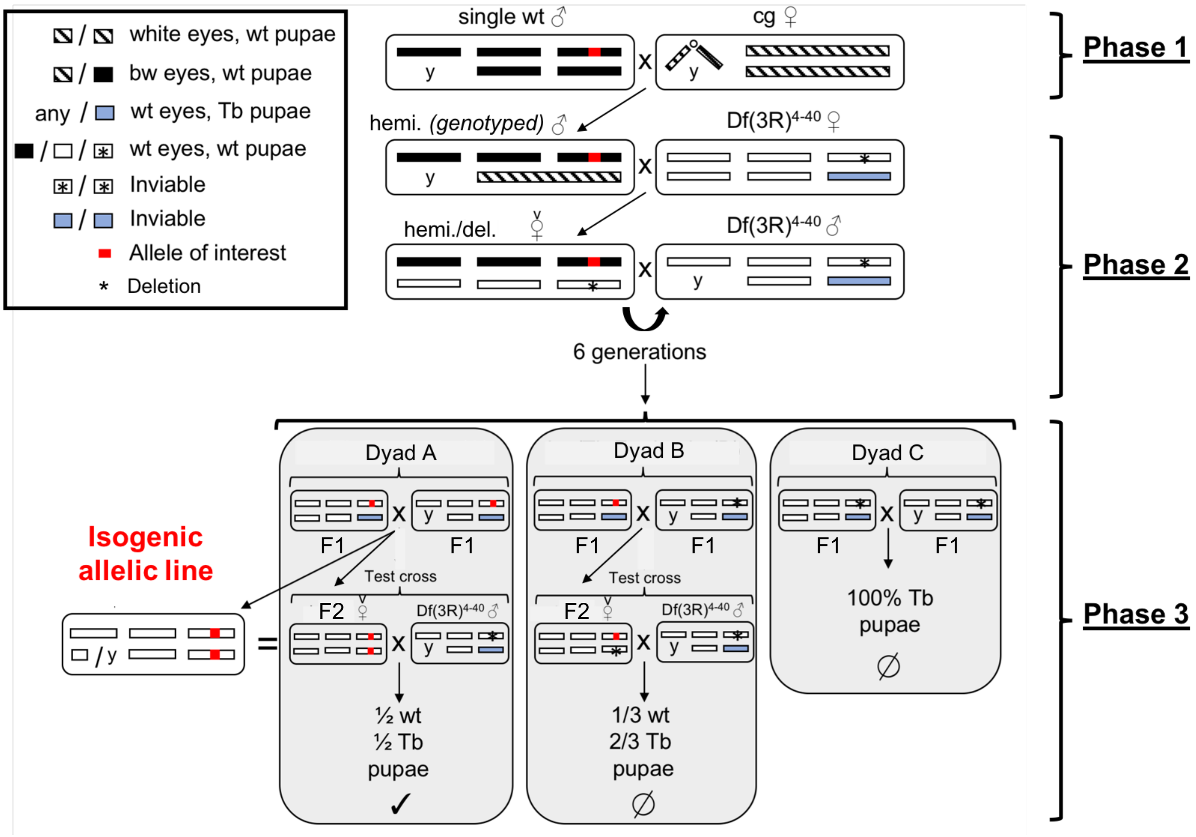

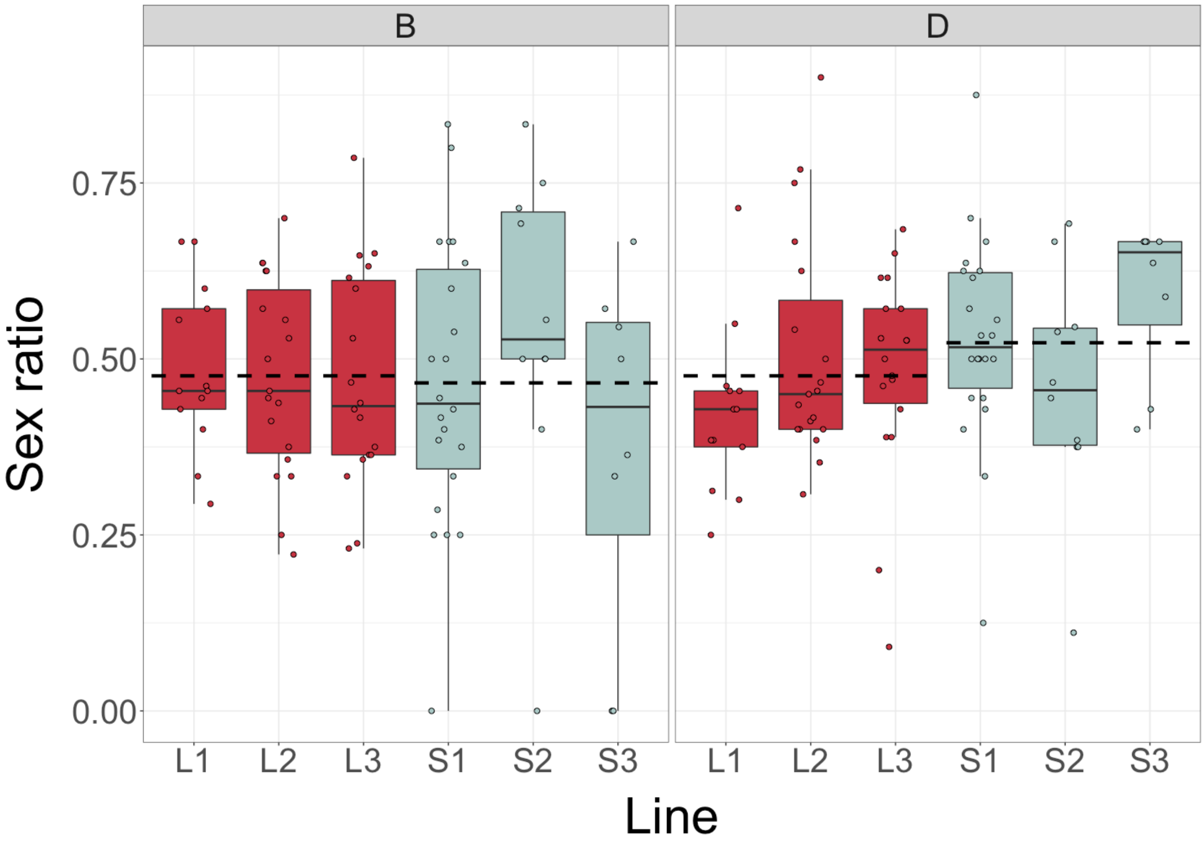


**Supplementary Figure 3.** Sex ratio among surviving offspring presented for each line (L1-3 and S1-3) and chromosomal complement (B and D). Allelic means represented by dashed lines. Sex ratio is defined as the proportion of males among offspring at eclosion.

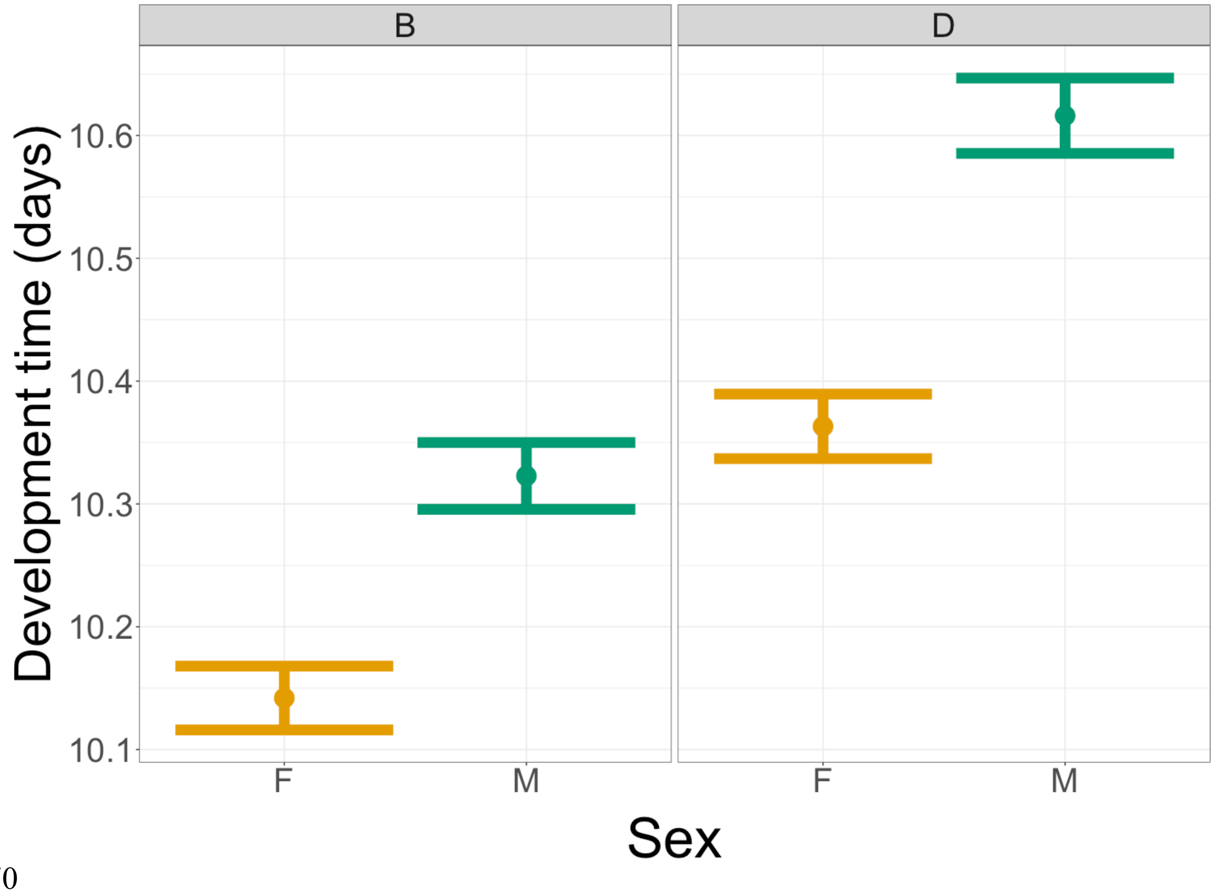


**Supplementary Figure 4.** Development time (days ±standard error) of each sex (Females(F)= yellow; males(M)= green), for each chromosomal complement (B and D).

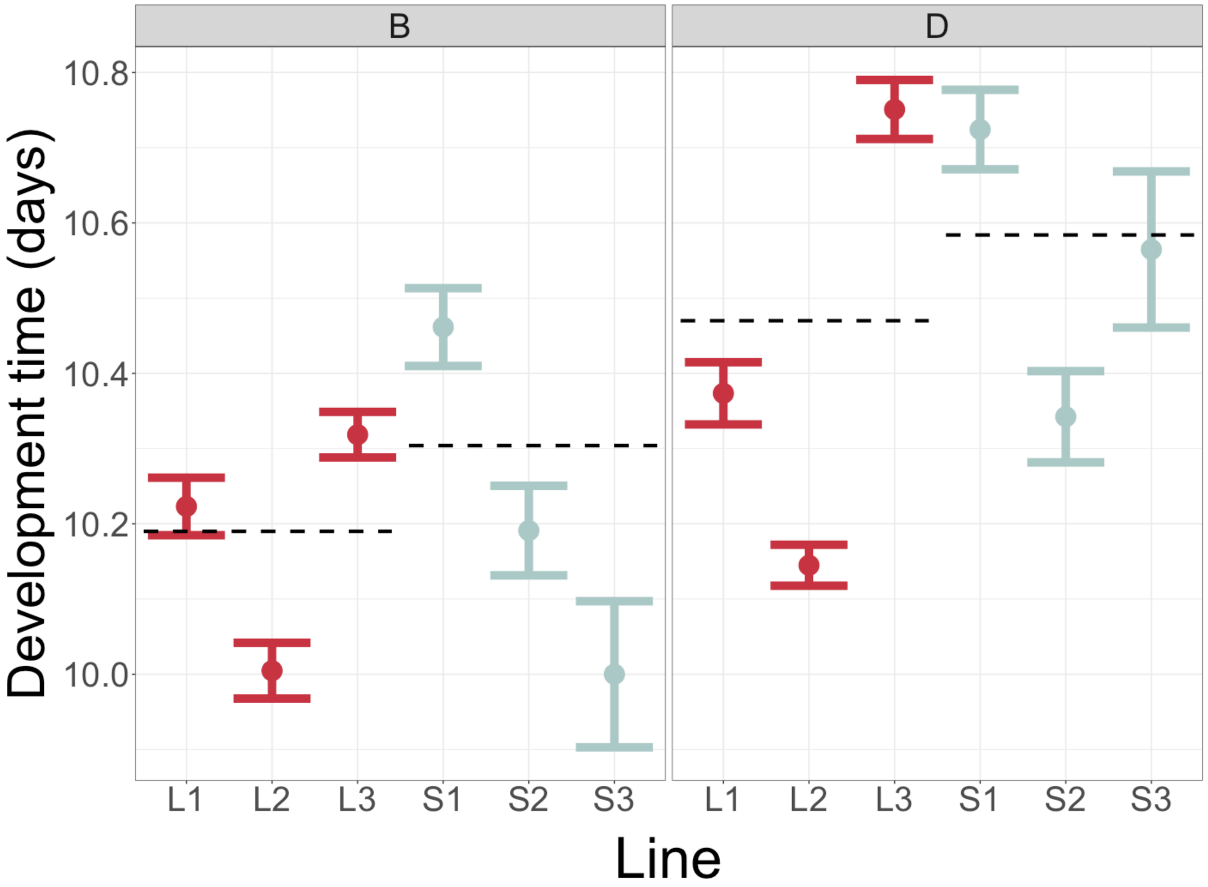


**Supplementary Figure 5.** Development time (days ±standard error) of *fru* allelic lines (L1-3 and S1-3), for each chromosomal complement (B and D). Allelic means represented by dashed lines.

**Supplementary Table 1. Results from Cox Proportional Hazard (CPH) models applied to lifespan data.** Five models were used. One was for all flies and then the data was split to have separate models for each chromosome complement (B and D) and sex (female or male). The first column indicates the set of data the model is applied to, while the second column indicates the term being tested in that model. CPH models use one level of a term as the reference level with a value of one. Other levels are then compared to this. The comparison made is shown in brackets as: (compared level:reference). Each term in a model has a hazard-ratio (H-R), a 95% confidence interval and a H-R p-value, which indicates if the compared level differs from the reference level. Also presented are $\chi_1^2$ and its p-value, indicating the contribution of each term to the overall risk of mortality.

| Model | Term (comparison) | HR | 95%-CI | HR p-value | $\chi_1^2$ | p-value |
|---|---|---|---|---|---|---|
| All flies | *fru* allele (S:L) | 1.318 | 1.126-1.544 | <0.001 | 0.139 | 0.71 |
| | Complement (D:B) | 0.519 | 0.44-0.612 | <0.001 | 43.79 | <0.001 |
| | Sex (M:F) | 0.531 | 0.449-0.627 | <0.001 | 31.886 | <0.001 |
| | Allele x complement (S/D:L/F) | 0.693 | 0.57-0.841 | <0.001 | 10.411 | 0.0013 |
| | Allele x sex (S/D:L/B) | 0.821 | 0.676-0.997 | 0.046 | 4.856 | 0.0276 |
| | Complement x sex (D/M:B/F) | 2.624 | 2.154-3.198 | <0.001 | 90.752 | <0.0001 |
| | Allele x complement x sex (S/D/M:L/B/F) | 1.258 | 0.852-1.856 | 0.249 | 1.331 | 0.249 |
| B only | *fru* allele (S:L) | 1.386 | 1.16-1.655 | <0.001 | 3.848 | 0.049 |
| | Sex (M:F) | 0.572 | 0.472-0.692 | <0.001 | 105.65 | <0.001 |
| | Allele x sex (S/D:L/B) | 0.731 | 0.561-0.953 | 0.02 | 5.368 | 0.021 |
| D only | *fru* allele (S:L) | 0.87 | 0.715-1.059 | 0.164 | 5.317 | 0.021 |
| | Sex (M:F) | 1.32 | 1.081-1.614 | 0.0066 | 10.705 | 0.001 |
| | Allele x sex (S/D:L/B) | 0.927 | 0.696-1.234 | 0.604 | 0.269 | 0.604 |
| Females only | *fru* allele (S:L) | 1.381 | 1.157-1.65 | <0.001 | 2.334 | 0.127 |
| | complement (D:B) | 0.542 | 0.449-0.655 | <0.001 | 14.879 | <0.001 |
| | Allele x complement (S/D:L/F) | 0.611 | 0.469-0.798 | <0.001 | 13.127 | <0.0001 |
| Males only | *fru* allele (S:L) | 1.039 | 0.854-1.263 | 0.705 | 1.276 | 0.259 |
| | complement (D:B) | 1.301 | 1.061-1.595 | 0.011 | 3.119 | 0.077 |
| | Allele x complement (S/D:L/F) | 0.772 | 0.58-1.029 | 0.077 | 3.117 | 0.077 |
